# Investigating the possible origin and transmission routes of SARS-CoV-2 genomes and variants of concern in Bangladesh

**DOI:** 10.1101/2021.05.24.444482

**Authors:** Abdullah Al Nahid, Ajit Ghosh

## Abstract

The COVID-19 pandemic induced by the SARS-CoV-2 virus and its variants has ravaged most countries around the world including Bangladesh. We have analyzed publicly available genomic data to understand the current COVID-19 outbreak scenario as well as the evolutionary origin and transmission routes of SARS-CoV-2 isolates in Bangladesh. All the early isolates as well as recent B.1.1.7 and B.1.351 variants had already spread across the major divisional cities of Bangladesh. A sex biasness towards male COVID-19 patient samples sequencing has observed over female in all age-group, that could be the trend in infection rate. Phylogenetic analysis indicated a total of 13 estimated countries, including Italy, India, United Kingdom, Saudi Arabia, United Arab Emirates, Germany, Australia, New Zealand, South Africa, Democratic Republic of the Congo, United States, Russia, and Denmark, could be the possible origin introduced SARS-CoV-2 isolates in Bangladesh due to regional and intercontinental travel. Recent, B.1.1.7 variant could be imported from a total of 7 estimated countries including UK, India, Nigeria, Spain, Ireland, Australia, and Indonesia, while South Africa and the United States are the most likely sources of B.1351 variant in Bangladesh. Based on these findings, public health strategies could be designed and implemented to reduce the local transmission of the virus.

## 1. Introduction

The world has been going through a global pandemic of acute respiratory disease named the coronavirus disease 2019 (COVID-19), caused by the severe acute respiratory syndrome coronavirus 2 (SARS-CoV-2). SARS-CoV-2 is a novel strain of *betacoronavirus* within the Coronaviridae family^1^, that was first reported from Wuhan, China, in December 2019^2^. SARS-CoV-2 possesses an enveloped single-stranded, positive-sense ~30kb long ribonucleic acid (RNA) genome^3^. The longest segment of the genome at 5’ end encodes for orf1ab polyprotein while the rest of the genome consists of genes encoding four structural proteins including the spike glycoprotein (S), envelope protein (E), membrane protein (M), and nucleocapsid protein (N)^4,5^. Since the identification of SARS-CoV-2, it has rapidly spread to 220 countries and territories including Bangladesh to date (https://www.worldometers.info/coronavirus), posing an unprecedented public health threat with over 165 million recorded cases of COVID-19, and more than 3.4 million deaths attributable to the disease worldwide as of May 20, 2021^6^. Bangladesh documented its first case of COVID-19 on March 8, 2020^7^. Since then, there have been 786,698 confirmed cases of the disease in the country, as well as 12,310 deaths with 1.56% case fatality rate (CFR)^6^.

Over the devastating course of the SARS-CoV-2 pandemic, several variants have appeared in different regions of the world. Of them, a more infectious variant with D614G mutation in the spike glycoprotein quickly became dominant after emergence, owing to the increased human-to-human transmission efficiency, amongst the globally circulating strains during the early stages of the pandemic^8–11^. According to a recent study, 98% of Bangladeshi SARS-CoV-2 isolates are of this common variant^12^. In addition to the pre-existing variants, several new variants of concern (VOCs) including B.1.1.7 or 501Y.V1, B.1.351 or 501Y.V2 and P.1 or 501Y.V3, have emerged independently from the United Kingdom (UK), South Africa, and Brazil respectively^13–15^. Out of these three VOCs, presence of B.1.1.7 and B.1.351 variants had been reported in Bangladesh (https://virological.org/t/detection-of-the-b-1-1-7-and-b-1-351-sars-cov-2-variants-in-bangladesh/668), as well as in dozens of other countries around the globe as of April 9, 2021 (https://cov-lineages.org/global_report.html). Although much is still unknown about these VOCs, preliminary epidemiological and phylogenetic studies suggest the B.1.1.7 variant is 43 to 90% (95% CI 38–130) more transmissible than pre-existing variants^13^, with a 50 to 100% higher reproduction number^16^, and increased mortality^17^. Furthermore, findings from other preliminary studies indicate the B1.351 variant is associated with higher viral load with potential advantage of enhanced transmissibility or immune escape^14,18–20^. This variant may also amplify the risk of infection in people who have already been immunized^21^.

As COVID-19 continues to wreak havoc due to rapid transformation of SARS-CoV-2, Bangladesh has begun a nationwide inoculation drive in order to restrain the virus, and administered more than 9.6 million doses of the Oxford-AstraZeneca vaccine to date (https://github.com/owid/covid-19-data/blob/master/public/data/vaccinations/country_data/Bangladesh.csv). However, the daily COVID-19 cases and deaths in the country have seen a steep rise recently^6^, which is found to be correlated with the increasing detection of B.1.351 variant circulating in the country^22^. Another genomic variant surveillance in Bangladesh, conducted from January 1 - March 24, 2021, observed a maximum 52% frequency of B.1.1.7 variant up until the second week of March 2021, and a dramatic rise in B.1.351 variant frequency of up to 81% in subsequent weeks (https://www.icddrb.org/news-and-events/news?id=874). In order to identify such emerging variants in the country, and monitor the viral evolution on a genomic level, scientists from Bangladesh have sequenced over 1,500 SARS-CoV-2 genomes, and deposited the sequences in the Global Initiative on Sharing All Influenza Data (GISAID) database^23^. Analyses of these genomic data may aid in understanding the genetical and evolutionary features of the virus^24^, as well as in evaluating the outcome of various disease control strategies in Bangladesh, ranging from quarantine measures to travel restrictions, both locally and internationally, to reduce the transmission rate of evolving lineages.

An epidemiological analysis on genome samples from Bangladesh revealed regional sources of the isolates including variants rather than country-specific origin information (https://nextstrain.org/community/CHRF-Genomics/ncovBangladesh@main). Another phylogenetic study on 64 Bangladeshi genome samples reported Germany, United Kingdom, India, and Saudi Arabia as the major possible sources of the studied isolates with no report on the likely origin of the variants^25^. Moreover, Nextstrain^26^ considered insufficient samples from Bangladesh for the global phylogenetics build of SARS-CoV-2 (https://nextstrain.org/ncov/global). Due to inadequate sampling of genome sequences from Bangladesh and lack of country-specific report on variant origin, most of these findings were not conclusive and fully reflective of the whole country. Thus, the recent rise in these VOCs necessitates a comprehensive genomic investigation of adequate and diverse samples to determine the country-specific transmission origin of Bangladeshi isolates including variants. As a result, this may help establish a firm understanding of viral epidemiology in the country, which is crucial for vaccine efficacy, clinical severity of infection, disease mortality, diagnostics and therapeutics, and public health safety precautions.

In this study, we aimed to identify the possible origin and transmission routes of SARS-CoV-2 isolates into Bangladesh, during the initial days of the pandemic, as well as for both B.1.1.7 and B.1.351 variants in recent times. We analyzed publicly available genomic data from countries of concern across all regions and estimated 3 separate time-resolved phylogenies with a total of 4,157 subsampled genome sequences. By utilizing the genomic epidemiology approach to the data, we demonstrate that both the initial viral outbreak and the recent upsurge of VOCs in Bangladesh was the result of several independent introductions of the virus linked to international travel.

## 2. Materials and methods

### 2.1 Data Acquisition & Filtering

A total of 97,040 unique SARS-CoV-2 genome sequences and metadata were downloaded from the GISAID database^23^ and subsequently divided into 4 subsets (**Table 1**) for downstream analyses. The G1 subset contained a total of 1,009 complete, human-host genomes from Bangladesh as of April 9, 2021 (**Dataset S1**). From the subset G1, a set of 612 high-coverage genomes till February 12, 2021 were considered for initial phylogeny and added to the G2 subset (**Dataset S2**). In addition, 86,314 complete genomes with a sample collection date prior to 1st May 2020 were included to the G2 subset from 48 additional countries and territories with higher COVID-19 prevalence during the early stages of the outbreak. For the G3 and G4 subsets, a total of 7,507 and 3,196 SARS-CoV-2 samples, including both B.1.1.7 and B.1.351 variants of 20I/501Y.V1 and 20H/501Y.V2 clade, respectively, from 101 and 31 different countries and territories around the world with reported cases of the virus and studied variants, were selected for variant-focused phylogenies. 5,100 of all G3 samples including 17 Bangladeshi samples (**Dataset S3**) belonged to the 20I/501Y.V1 clade, while 2,255 of all G4 samples along with 47 samples from Bangladesh (**Dataset S4**) belonged to the 20H/501Y.V2 clade. Sample submission cutoff date similar to G1 subset had been applied to both G3 and G4 subsets. Additional daily COVID-19 confirmed case related data of Bangladesh were downloaded from the World Health Organization (WHO) website (https://covid19.who.int/WHO-COVID-19-global-data.csv). Sample metadata of all subsets were checked in terms of data availability and any sample with incomplete collection date were excluded. Prior to phylogenetic analysis, G2, G3 and G4 subset samples were quality filtered and genome sequences with ambiguous characters were omitted.

**Table 1:** Summary of Collected Samples in Subsets

| Subset | Total Countries and Territories | Sample Time Period | Total Samples | Bangladeshi Samples |
| --- | --- | --- | --- | --- |
| G1 | 1 | Submission till April 9, 2021 | 1,009 | 1,009 |
| G2 | 49 | **For Bangladeshi samples:**<br>Submission till February 12, 2021<br>**For rest of the samples:**<br>Collection till April 30, 2020 | 86,926 | 612 |
| G3 | 101 | Submission till April 9, 2021 | 7,507 | 17 |
| G4 | 31 | Submission till April 9, 2021 | 3,196 | 47 |

### 2.2 Exploratory Data Analysis

The metadata of G1 subset samples was analyzed with Python version 3.8.5^27^ to investigate the Nextstrain clade distribution among divisions, as well as the gender and age-group distribution in Bangladeshi samples. G1 samples with no division data were labeled NA (Not Available), and the missing gender and age-group data were discarded before plotting with ggplot2 version 3.3.328 and tidyverse version 1.3.0^29^ packages of R programming language version 3.6.3^30^.

### 2.3 Phylogenetic Analysis

Quality filtered sequences of G2 subset were aligned to the reference SARS-CoV-2 sequence (NC_045512.2)^31^ using MAFFT version 7.475^32^. The aligned genomes were subsampled afterwards. All 612 genomes from Bangladesh within the G2 subset were assigned as focal samples during the subsampling process. In addition, 10 samples per month were chosen from 48 other countries and territories available in the subset based on genetic similarity to focal samples. A total of 1,966 subsampled genomes were used to construct a maximum-likelihood phylogenetic tree with IQ-TREE version 2.0.3^33^, and further refined by TreeTime version 0.8.1^34^. The complete analysis pipeline is available online (https://github.com/nextstrain/ncov, version 3). For visualization, auspice version 0.5.0 (https://auspice.us) by Nextstrain^26^ was used. Analyses performed on G2 subset samples were also performed on both G3 and G4 subsets to build phylogenetic trees for B.1.1.7 and B.1.351 variants using 1,546 and 645 subsampled genomes, respectively. During the subsampling process of both subsets, 5 samples a month were selected for each country present in the corresponding subset except Bangladesh. The interactive phylogenetic builds are available online at: https://github.com/nahid18/ncov-bd.

## 3. Results

### 3.1 Data Exploration for the current scenario of COVID-19 cases in Bangladesh

To better understand the current COVID-19 situation in Bangladesh, the pattern of daily reported COVID-19 cases was analyzed. The highest peak of nearly 8,000 confirmed cases per day was recently reported in the early April 2021 (**Fig. 1A**). The metadata of 1009 complete human-host SARS-CoV-2 genomes from Bangladesh (**Dataset S1**) accessible via GISAID^23^ were examined. The overall Nextstrain clade distribution across all the major divisional cities of Bangladesh (**Fig. 1B**), reveals that samples from Dhaka and Sylhet divisions contain the 20I/501Y.V1 clade and primarily start clustering around January 2021. Furthermore, samples from the Dhaka, Chattogram, Khulna, and Mymensingh divisions include the 20H/501Y.V2 clade that has been the most prevalent during March to April 2021. This raises the alarming possibility that these VOCs are now dispersed in Bangladesh. We have also looked at the gender and age-group distribution within the available samples and found out that samples from male patients are more frequently sequenced in almost every age-group except 10-19, than female patients (**Fig. 1C** and **Supplementary Table S4**).

**Figure 1A:**
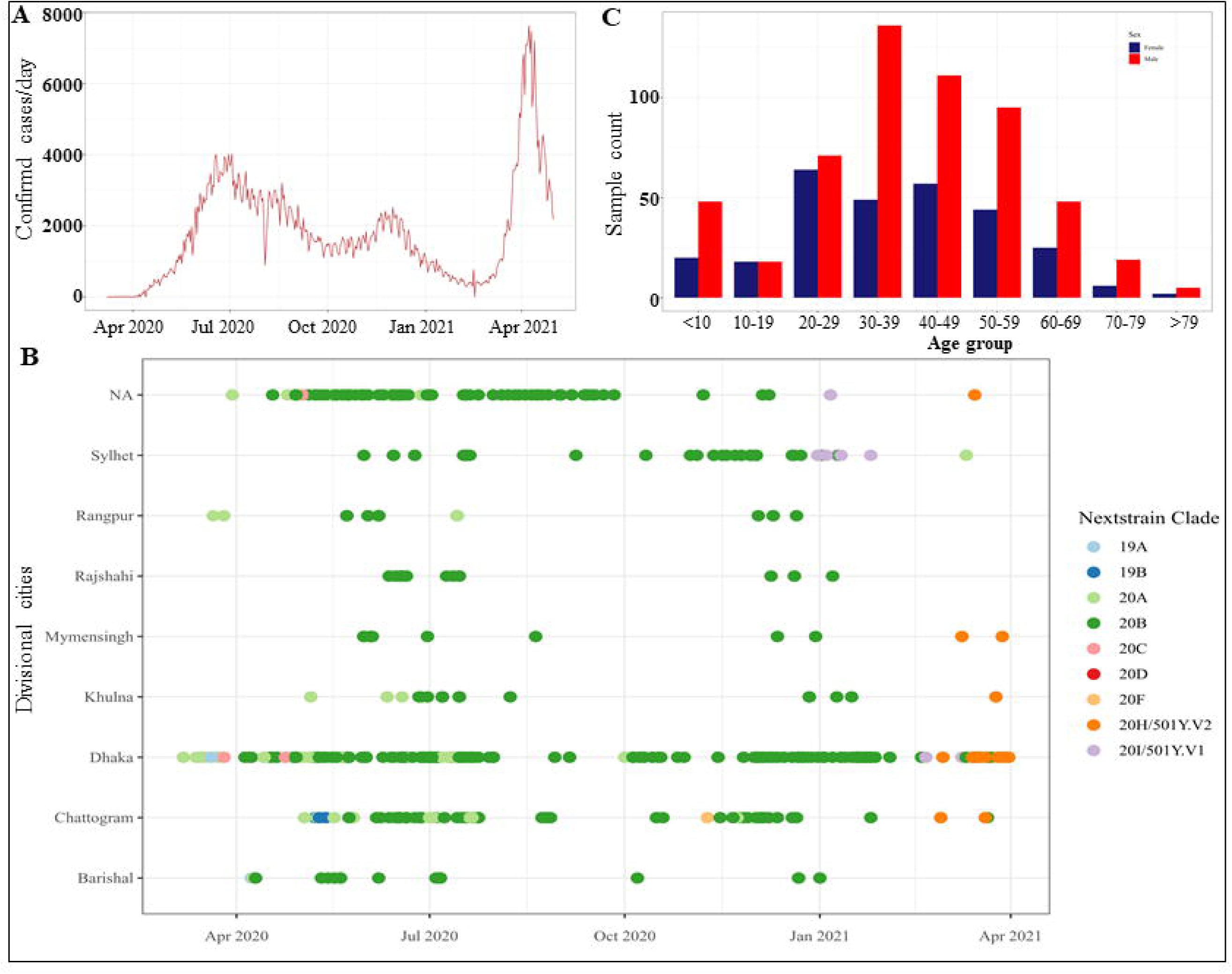
Trend of COVID-19 confirmed cases in Bangladesh over the course of the pandemic. **Figure 1B:** Distribution of Nextstrain clades and Bangladeshi Divisions in SARS-CoV-2 samples isolated in Bangladesh (n=1009). NA represents the samples with unassigned division data. **Figure 1C:** Distribution of sex and age-group in Bangladeshi SARS-CoV-2 samples (n=836). The male and female samples are represented via red and dark-blue colors, respectively.

### 3.2 Phylogenetic analysis of SARS-CoV-2 isolates of Bangladesh

An independent phylogenetic analysis has been performed using a curated collection of 86,926 SARS-CoV-2 genome sequences covering all regions and most countries/territories where the COVID-19 infection rate was high during the early stages of the pandemic to precisely trace the origin of initial SARS-CoV-2 isolates of Bangladesh. The generated phylogenetic tree containing a total of 1,966 subsampled genomes, showed that all 612 high-coverage genome sequences from Bangladesh are scattered across the tree (**Fig. 2**), suggesting multiple introductions of the virus into the country, as well as grouped in several distinct clusters indicating community transmission. The two well-defined clusters of Bangladeshi samples within 20B clade contain a total of 482 descendants and share common ancestry with sequences isolated in Italy (EPI_ISL_469050, EPI_ISL_494761, EPI_ISL_494774). In addition, multiple instances of Bangladeshi samples with an estimated foreign origin have been found in 5 separate clades distributed within the tree (**Supplementary Table S5**). In detail, Bangladeshi sample with GISAID accession no. EPI_ISL_458133 is close to the sample EPI_ISL_509705 from the United States within 19A clade. Two samples from Bangladesh, EPI_ISL_450339 and EPI_ISL_450341, have high similarity with the Australian EPI_ISL_410718 sample of 19B clade. Another Bangladeshi sample EPI_ISL_450345 within 19B clade is showing high similarity with an Indian sample EPI_ISL_450324.

**Figure 2:**
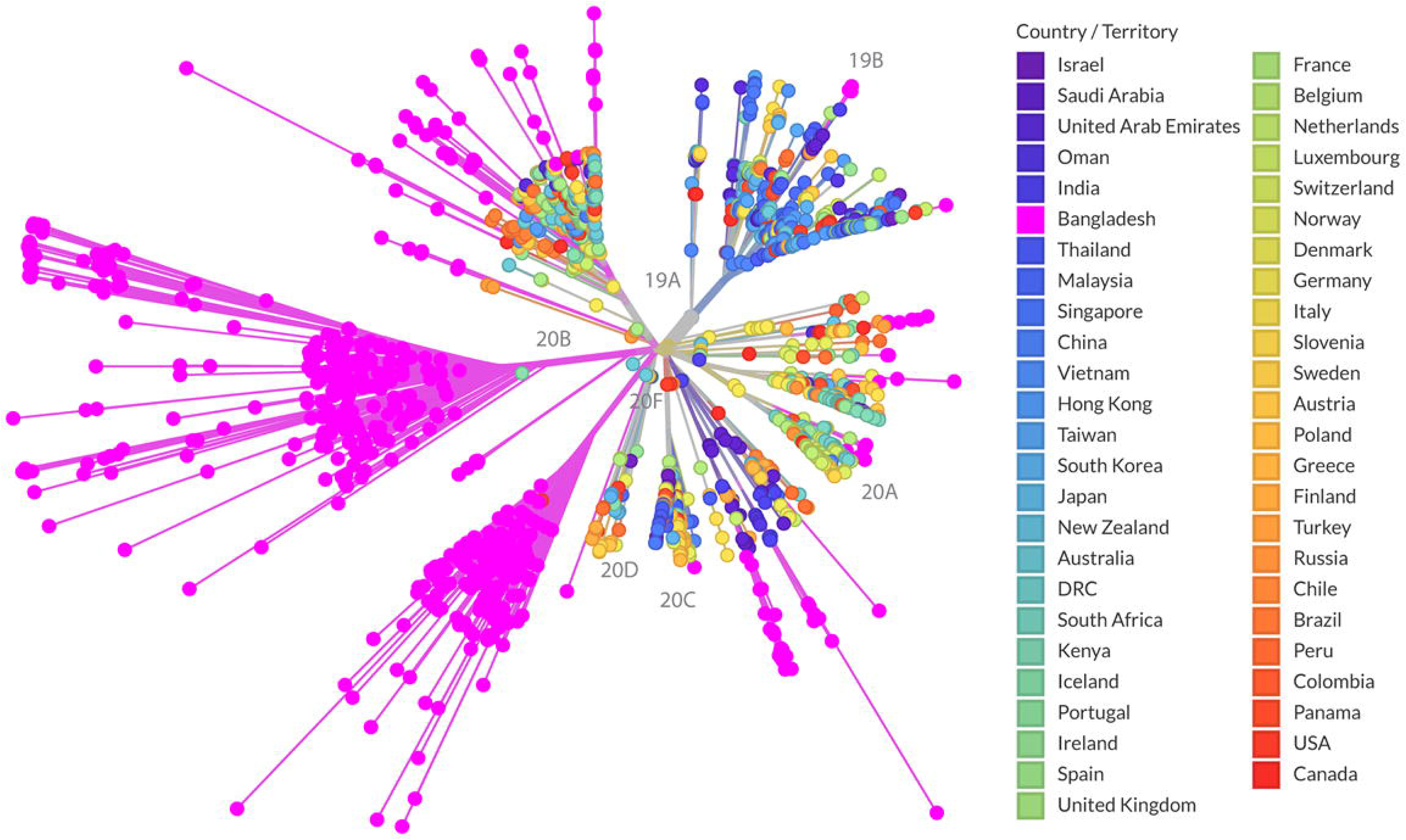
Phylogenetic tree of initial SARS-CoV-2 isolates in an unrooted representation. Out of 86,926 analyzed genomes, the tree was constructed using a total of 1,966 subsampled genomes and visualized using auspice version 0.5.0 (https://auspice.us). Pink dots represent all 612 Bangladeshi samples present in the phylogeny.

In addition to the large clusters of Bangladeshi samples, several clusters of few samples are identified as well throughout the tree, indicating the local transmission on a smaller scale. Within the 20A clade, EPI_ISL_445244 and 3 other samples from Bangladesh are close to Italian EPI_ISL_412973 and EPI_ISL_451306 samples. A group of 10 Bangladeshi samples including EPI_ISL_774877, EPI_ISL_605887 and EPI_ISL_774896 within the same clade have a shared ancestry with the EPI_ISL_707964 sample isolated in Germany. Another group of 11 Bangladeshi clustered as 20A, including EPI_ISL_774900 and EPI_ISL_605910, are similar to the Russian EPI_ISL_450288 sample. Similarly, a total of 19 Bangladeshi 20A samples including EPI_ISL_466637 and EPI_ISL_450343 have close similarity to the EPI_ISL_513222 sample from Saudi Arabia. In addition, four samples within 20A clade from Bangladesh including EPI_ISL_468070 and EPI_ISL_774893 are highly related to the sample EPI_ISL_429255 from the Democratic Republic of the Congo (DRC). Indian EPI_ISL_435065, and South African EPI_ISL_450299 sample have close resemblance to the Bangladeshi EPI_ISL_477134 and EPI_ISL_605889 samples of the 20A clade, respectively.

Although most of the 20B samples from Bangladesh are found to have Italian ancestry, samples from the United Kingdom, United Arab Emirates and New Zealand are also showing high similarity to Bangladeshi clustered detected across the 20B clade. EPI_ISL_477130, EPI_ISL_774889, EPI_ISL_774890 and 5 other samples from Bangladesh are closely related to the EPI_ISL_488766 sample from the United Kingdom. A total of 10 samples from Bangladesh including EPI_ISL_483630 and EPI_ISL_464162 are close to the EPI_ISL_579166 sample isolated in New Zealand. Moreover, a combined set of 8 Bangladeshi samples of 20B clade in 3 separate groups are closely related to 3 samples with GISAID accession no. EPI_ISL_520738, EPI_ISL_520717 and EPI_ISL_528686 from the United Arab Emirates (UAE). In 20C clade, Bangladeshi EPI_ISL_477128 sample is highly similar to the EPI_ISL_614672 and EPI_ISL_614674 samples from Denmark.

Based on the evidence indicated by the phylogenetic tree, it is evident that SARS-CoV-2 isolates have been introduced into Bangladesh from several countries of all regions, except South America, during the initial outbreak of COVID-19. Multiple introductions from a single country such as Italy, India and UAE have occurred as well.

### 3.3 Phylogenetic analysis of B.1.1.7 variant samples in Bangladesh as compared to other countries

A time-resolved phylogenetic tree was created containing 1,546 subsampled genomes including 641 samples of B.1.1.7 variant to develop a better understanding of the evolutionary relationship and potential propagation patterns of this variant in Bangladesh. All 17 Bangladeshi samples are found scattered across the 20I/501Y.V1 clade within the tree (**Fig. 3**), suggesting multiple viral introductions into Bangladesh from several countries (**Supplementary Table S6**). The sample EPI_ISL_1360439 from Bangladesh is closely related to the sample EPI_ISL_756846 isolated in the United Kingdom. Another Bangladeshi sample EPI_ISL_906098 is highly similar to a sample EPI_ISL_837380 from Ireland. Two samples-EPI_ISL_1508943 and EPI_ISL_1508946, are close to the EPI_ISL_1118931 sample from Indonesia. A total of 6 samples including EPI_ISL_1509000 and EPI_ISL_1360451 are related to the Nigerian EPI_ISL_985080, EPI_ISL_872625 samples. Three distinct Bangladeshi samples (EPI_ISL_1498132, EPI_ISL_1360445 and EPI_ISL_906091) have shown similarity with samples from India (EPI_ISL_995709), Australia (EPI_ISL_812441) and Spain (EPI_ISL_1059975), respectively.

**Figure 3:**
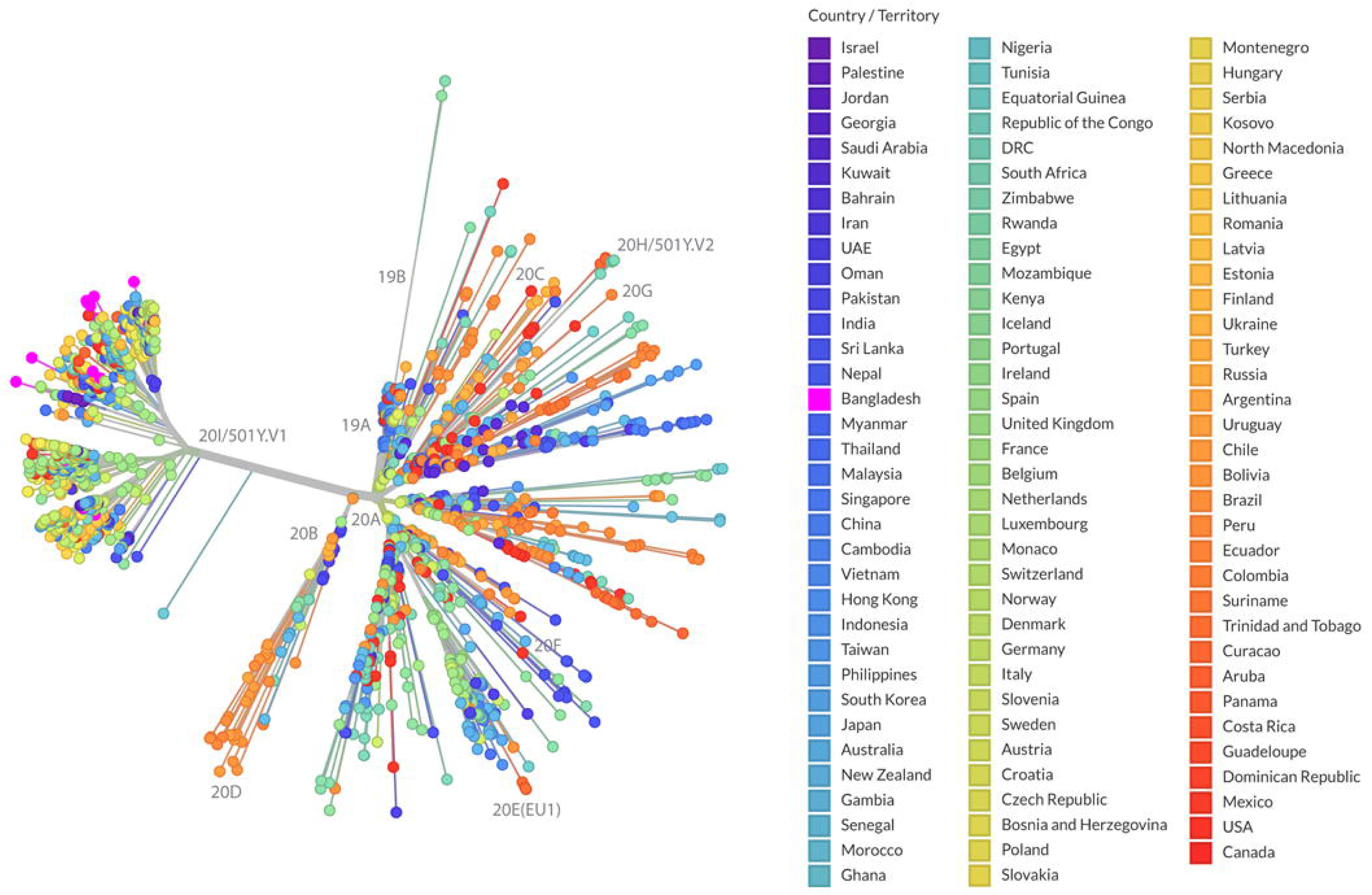
Phylogenetic tree of B.1.1.7 variant samples in an unrooted representation. Out of 7,507 analyzed B.1.1.7 variant samples, the tree was constructed using a total of 1,546 subsampled genomes and visualized using auspice version 0.5.0 (https://auspice.us). All 17 analyzed B.1.1.7 variant samples of Bangladesh are represented by the pink colored dots.

### 3.4 Phylogenetic analysis of B.1.351 variant Bangladeshi samples in a global point of view

In order to identify the origin of the B.1.351 variant isolates in Bangladesh, we analyzed 3,196 samples from 31 countries and territories, the bulk of which were B.1.351 variant. The resulting phylogenetic tree (**Fig. 4**) generated from the 645 subsampled sequences, reveals 46 out of 47 B.1351 variant samples of Bangladesh clustered under a single node within 20H/501Y.V2 clade, manifesting clear signs of local transmission with single ancestral source, with paternal node being shared with samples isolated in South Africa. Samples from France and Turkey are placed close to the Bangladeshi samples in the phylogenetic tree. The remaining lone sample (EPI_ISL_1508892) that did not cluster with other samples had the closest ancestral genotype of United States origin (EPI_ISL_1097839). Based on the data (**Supplementary Table S7**), it is highly likely that South Africa and the United States are the estimated origins to have introduced the B.1.351 variant to Bangladesh.

**Figure 4:**
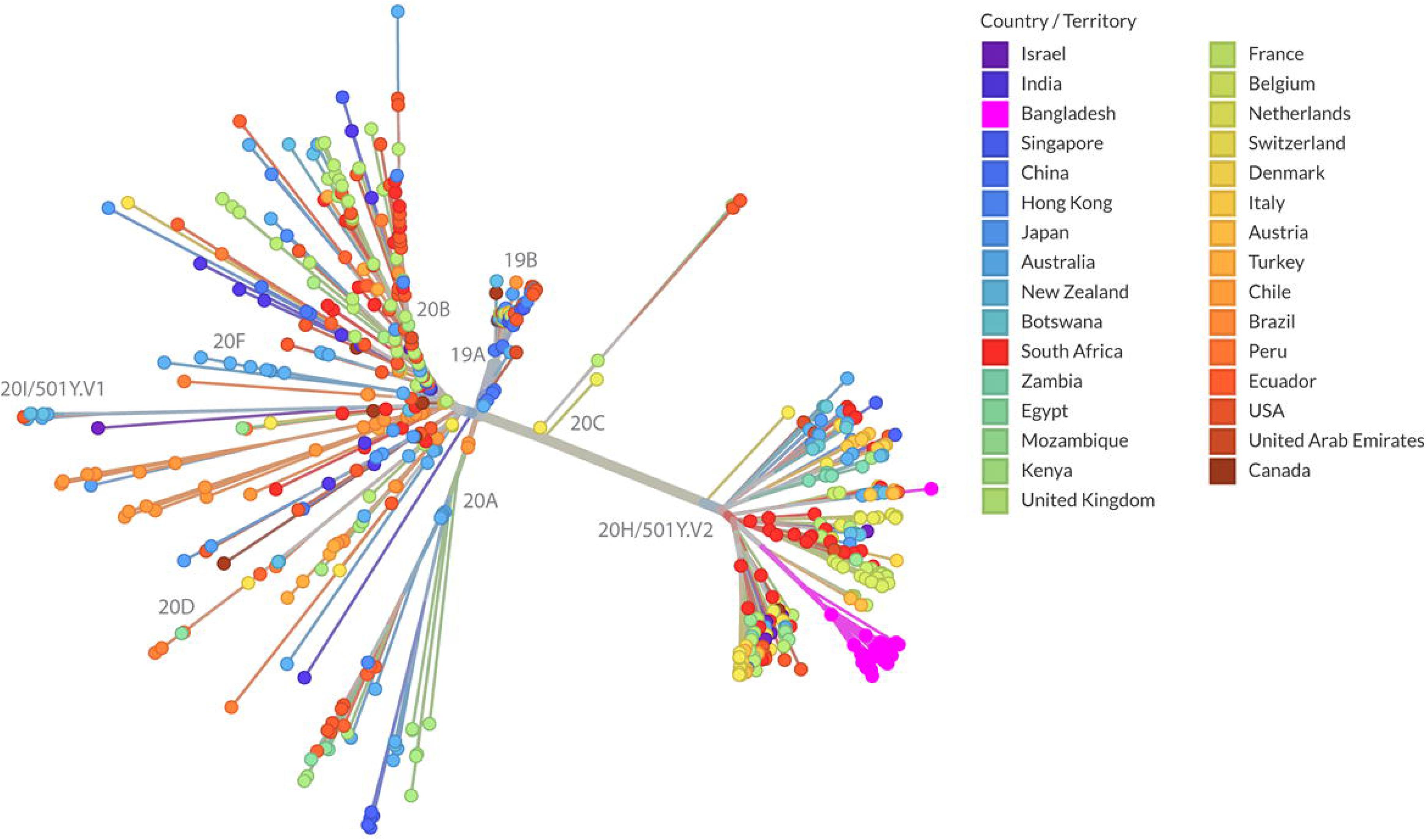
Phylogenetic tree of B.1.351 variant samples in an unrooted representation. Out of 3,196 analyzed B.1.351 variant samples, the tree was constructed using a total of 645 subsampled genomes and visualized using auspice version 0.5.0 (https://auspice.us). Pink colored dots are the representatives of all 47 B.1.351 variant samples of Bangladesh.

## Discussions

As the global pandemic of COVID-19 progresses, new SARS-CoV-2 variants that are highly pathogenic and possibly more transmissible in nature than pre-existing variants continues to emerge due to mutational changes in the viral genome, causing a second wave of COVID-19 in several regions and countries around the world^35,36^. From the COVID-19 confirmed cases data, a severe second wave is detected in Bangladesh from late March to early April 2021 (**Fig. 1A**) as compared to the first wave of April to August 2020^7^. The exploratory analysis on Bangladeshi genomic sequences reveals a high prevalence of B.1.351 variants could be a major determining factor for the second wave of COVID-19 in the country, which is consistent with an already published study^22^. Moreover, we have reported that both B.1.1.7 and B.1.351 variant samples in Bangladesh are now dispersed in several divisions of Bangladesh including Dhaka, Sylhet, Chattogram, Mymensingh, and Khulna (**Fig. 1B**). A continuous trend of male-female disparity in terms of sample sequencing in almost every age group was observed (**Fig. 1C**), with more male patient samples being sequenced than female patient samples. This is most likely attributed to the fact that male COVID-19 patients has a far higher infection and mortality rate than females, as several studies have reported similar gender-based differentiation of COVID-19 severity in Bangladesh^37,38^. Biological sex plays an important role in the COVID-19 infection outcome by manifesting the infection susceptibility, pathogenesis, innate and adaptive immune responses, level of inflammation, and capacity of tissue repair. Presence of a more robust T cell activation mechanism and higher levels of innate immune cytokines in female as compared to male patients, could be the possible explanation for this kind of gender biasness in COVID-19^39^. A male dominance in COVID-19 mortality rate has been consistent with the other viral pathogenesis^37^.

Phylogenetic analyses on all the subsets have helped us to unravel the estimated origins of SARS-CoV-2 isolates as well as both B.1.1.7 and B.1.351 variants in Bangladesh. Based on the evidence presented in the phylogenetic trees (**Fig. 2-4**), a total of 17 unique countries from all regions of the world, except South America, have collectively introduced initial SARS-CoV-2 isolates as well as two analyzed VOCs into Bangladesh (**Fig 5**). Of these countries, initial Bangladeshi samples are showing similarity with the sequences reported from Italy, India, UK, Saudi Arabia, UAE, Germany, Australia, New Zealand, South Africa, DRC, US, Russia, and Denmark. A subset of these countries was reported to be the estimated origins of SARS-CoV-2 in Bangladesh in few other studies^25,40^. Multiple introductions from a single country including Italy, India, UK, Saudi Arabia have occurred as well. Furthermore, B.1.1.7 variant samples of Bangladesh are found to have close similarity to the samples from UK, India, Nigeria, Spain, Ireland, Australia, and Indonesia. In case of B.1.351 variant, it is highly possible that South Africa and the USA are the estimated origins of this variant in Bangladesh. From these findings, It is also evident that multiple variants from several countries including UK, India, South Africa, US, and Australia have arrived in Bangladesh because of regional and intercontinental travel. Based on the widespread distribution of several distinct clusters throughout the phylogenetic tree, these foreign isolates might be transmitted locally in different areas of Bangladesh during the COVID-19 pandemic.

**Figure 5:**
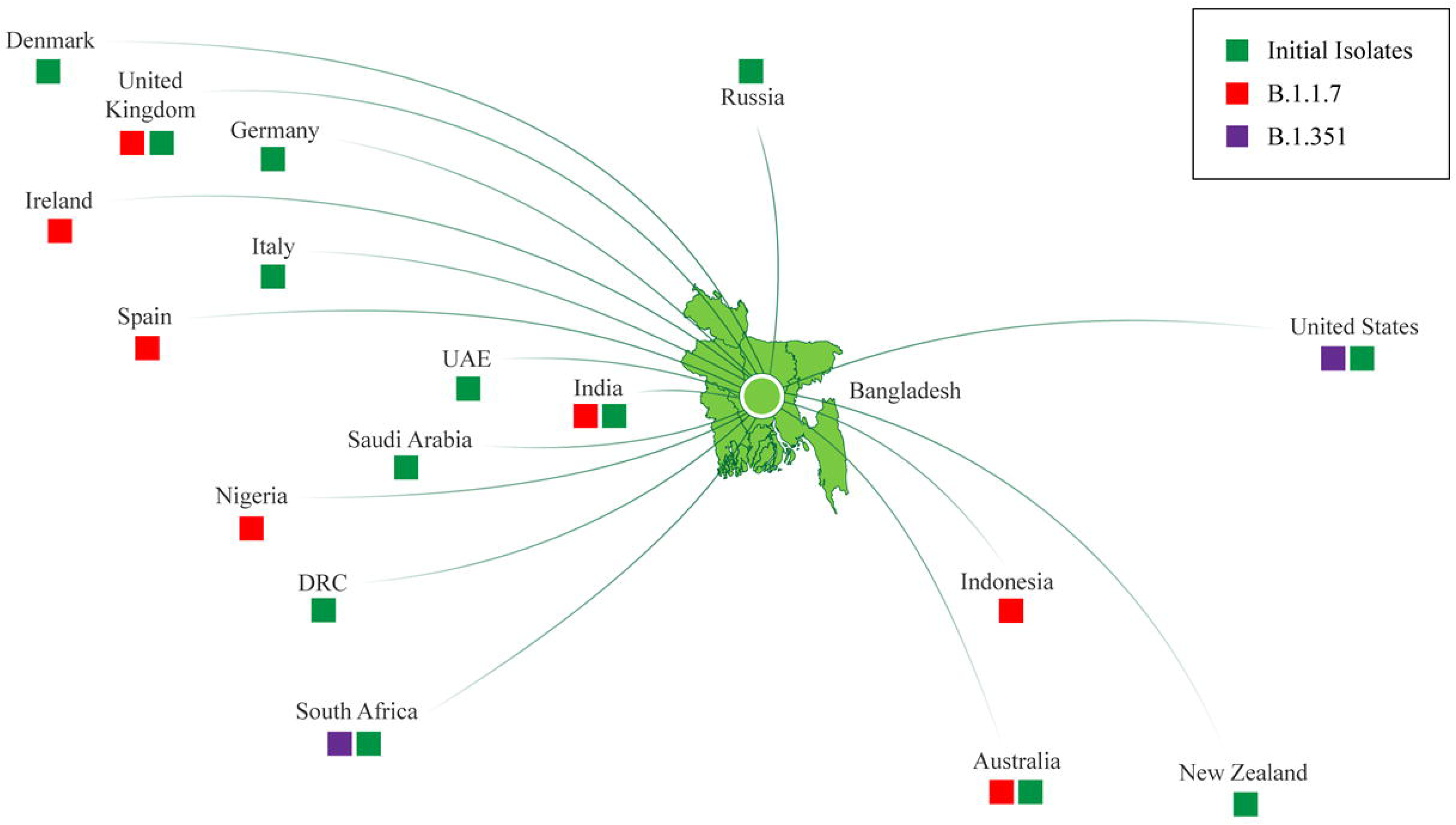
Estimated origins and possible transmission routes of early SARS-CoV-2 isolates and B.1.1.7, B.1.351 variants into Bangladesh.

Findings from this study should be taken into consideration within the context of a few caveats. During the subsampling process, a fixed number of samples were selected for the final grouping, hence may not be included all foreign samples that are possibly close to the focal samples. Besides, samples for the variant phylogenies may be biased towards the country of initial emergence of the variants. Despite these limitations, the findings regarding the origin of these variants are very alarming and should be brought to the notice of policymakers and public health authorities right away to prevent further escalation of viral transmission on a community level in Bangladesh.

## Conclusions

This study illustrates the contribution of both regional and intercontinental travel in the spread of SARS-CoV-2 VOCs in Bangladesh. Continuous genomic surveillance is crucial while the COVID-19 pandemic unfolds in order to closely monitor the ever-changing circulation pattern of SARS-CoV-2 variants in Bangladesh since these findings have major ramifications for mitigation and vaccination tactics. Given how quickly new variants can propagate across the world due to air travel without being detected for a long time, these findings are critical for administrative authorities and public health officials in terms of implementing effective interventions and guidelines including but not limited to travel restrictions and mandatory quarantine or lockdown measurements to prevent further proliferation of these variants in Bangladesh. Immunization efforts must also be continued and enforced to keep the infection rates down.

## Data availability

The authors declare that all the data will be available without any restrictions.

## Authors’ Contributions

AG conceived the idea and designed the experiments. AAN performed all the experiments, analyzed. AAN wrote the manuscript. All authors read the manuscript and approved the final version.

## Acknowledgements

Authors acknowledge the logistic support and laboratory facilities of the Department of Biochemistry and Molecular Biology, Shahjalal University of Science and Technology, Sylhet, Bangladesh.

## Supporting Information

**Table S1.** Number of samples for the initial phylogeny from different countries/ territories (n=86,926)

**Table S2.** Number of B.1.1.7 samples from different countries/ territories (n=7,507)

**Table S3.** Number of B.1.351 samples from different countries/ territories (n=3,196)

**Table S4.** Distribution of Age Group and Sex in G1 subset without missing values (n=836)

**Table S5.** Estimated origins of initial representative samples of Bangladesh

**Table S6.** Estimated origins of B.1.1.7 variant representative samples of Bangladesh

**Table S7.** Estimated origins of B.1.351 variant representative samples of Bangladesh

**Dataset S1.** Detail information of G1 subset containing a total of 1,009 genomes

**Dataset S1.** Detail information of G2 subset containing a total of 86,926 genomes

**Dataset S3.** Detail information of G3 subset containing a total of 7,507 genomes

**Dataset S4.** Detail information of G4 subset containing a total of 3,196 sequences

## References

1. Gorbalenya, A. E. et al. The species Severe acute respiratory syndrome-related coronavirus◻: classifying 2019-nCoV and naming it SARS-CoV-2. Nat. Microbiol. 5, 536–544 (2020).

2. Zhou, P. et al. A pneumonia outbreak associated with a new coronavirus of probable bat origin. Nature 579, 270–273 (2020).

3. Naqvi, A. A. T. et al. Insights into SARS-CoV-2 genome, structure, evolution, pathogenesis and therapies: Structural genomics approach. Biochim. Biophys. Acta BBA - Mol. Basis Dis. 1866, 165878 (2020).

4. Hu, B., Guo, H., Zhou, P. & Shi, Z.-L. Characteristics of SARS-CoV-2 and COVID-19. Nat. Rev. Microbiol. 19, 141–154 (2021).

5. Michel, C. J., Mayer, C., Poch, O. & Thompson, J. D. Characterization of accessory genes in coronavirus genomes. Virol. J. 17, 131 (2020).

6. Dong, E., Du, H. & Gardner, L. An interactive web-based dashboard to track COVID-19 in real time. Lancet Infect. Dis. 20, 533–534 (2020).

7. Imran, Md. A., Noor, I. U. & Ghosh, A. Impact of Lockdown Measures and Meteorological Parameters on the COVID-19 Incidence and Mortality Rate in Bangladesh. Infect. Microbes Dis. 3, 41–48 (2021).

8. Korber, B. et al. Tracking Changes in SARS-CoV-2 Spike: Evidence that D614G Increases Infectivity of the COVID-19 Virus. Cell 182, 812–827.e19 (2020).

9. Yurkovetskiy, L. et al. Structural and Functional Analysis of the D614G SARS-CoV-2 Spike Protein Variant. Cell 183, 739–751.e8 (2020).

10. Volz, E. et al. Evaluating the Effects of SARS-CoV-2 Spike Mutation D614G on Transmissibility and Pathogenicity. Cell 184, 64–75.e11 (2021).

11. Harrison, A. G., Lin, T. & Wang, P. Mechanisms of SARS-CoV-2 Transmission and Pathogenesis. Trends Immunol. 41, 1100–1115 (2020).

12. Hasan, Md. M. et al. Global and local mutations in Bangladeshi SARS-CoV-2 genomes. Virus Res. 297, 198390 (2021).

13. Davies, N. G. et al. Estimated transmissibility and impact of SARS-CoV-2 lineage B.1.1.7 in England. Science 372, (2021).

14. Tegally, H. et al. Detection of a SARS-CoV-2 variant of concern in South Africa. Nature 592, 438–443 (2021).

15. Faria, N. R. et al. Genomics and epidemiology of the P.1 SARS-CoV-2 lineage in Manaus, Brazil. Science (2021) doi:10.1126/science.abh2644.

16. Volz, E. et al. Assessing transmissibility of SARS-CoV-2 lineage B.1.1.7 in England. Nature 1–6 (2021) doi:10.1038/s41586-021-03470-x.

17. Davies, N. G. et al. Increased mortality in community-tested cases of SARS-CoV-2 lineage B.1.1.7. Nature 1–5 (2021) doi:10.1038/s41586-021-03426-1.

18. Cele, S. et al. Escape of SARS-CoV-2 501Y.V2 from neutralization by convalescent plasma. Nature 1–6 (2021) doi:10.1038/s41586-021-03471-w.

19. Wibmer, C. K. et al. SARS-CoV-2 501Y.V2 escapes neutralization by South African COVID-19 donor plasma. Nat. Med. 27, 622–625 (2021).

20. Pearson, C. A. B., Russell, T. W., Davies, N. & Kucharski, A. J. Estimates of severity and transmissibility of novel SARS-CoV-2 variant 501Y.V2 in South Africa. CMMID Repository https://cmmid.github.io/topics/covid19/sa-novel-variant.html (2021).

21. Planas, D. et al. Sensitivity of infectious SARS-CoV-2 B.1.1.7 and B.1.351 variants to neutralizing antibodies. Nat. Med. 1–8 (2021) doi:10.1038/s41591-021-01318-5.

22. Saha, S. et al. COVID-19 rise in Bangladesh correlates with increasing detection of B.1.351 variant. BMJ Glob. Health 6, e006012 (2021).

23. Shu, Y. & McCauley, J. GISAID: Global initiative on sharing all influenza data – from vision to reality. Eurosurveillance 22, (2017).

24. Siddiqe, R. & Ghosh, A. Genome-wide in silico identification and characterization of Simple Sequence Repeats in diverse completed SARS-CoV-2 genomes. Gene Rep. 23, 101020 (2021).

25. Shishir, T. A., Naser, I. B. & Faruque, S. M. In silico comparative genomics of SARS-CoV-2 to determine the source and diversity of the pathogen in Bangladesh. PLOS ONE 16, e0245584 (2021).

26. Hadfield, J. et al. Nextstrain: real-time tracking of pathogen evolution. Bioinformatics 34, 4121–4123 (2018).

27. Van Rossum, G. & Drake, F. L. Python 3 Reference Manual. (CreateSpace, 2009).

28. Wickham, H. ggplot2: Elegant Graphics for Data Analysis. (Springer-Verlag New York, 2016).

29. Wickham, H. et al. Welcome to the tidyverse. J. Open Source Softw. 4, 1686 (2019).

30. R Core Team. R: A Language and Environment for Statistical Computing. (R Foundation for Statistical Computing, 2013).

31. Wu, F. et al. A new coronavirus associated with human respiratory disease in China. Nature 579, 265–269 (2020).

32. Katoh, K. & Standley, D. M. MAFFT Multiple Sequence Alignment Software Version 7: Improvements in Performance and Usability. Mol. Biol. Evol. 30, 772–780 (2013).

33. Minh, B. Q. et al. IQ-TREE 2: New Models and Efficient Methods for Phylogenetic Inference in the Genomic Era. Mol. Biol. Evol. 37, 1530–1534 (2020).

34. Sagulenko, P., Puller, V. & Neher, R. A. TreeTime: Maximum-likelihood phylodynamic analysis. Virus Evol. 4, (2018).

35. Cacciapaglia, G., Cot, C. & Sannino, F. Second wave COVID-19 pandemics in Europe: a temporal playbook. Sci. Rep. 10, 15514 (2020).

36. Salyer, S. J. et al. The first and second waves of the COVID-19 pandemic in Africa: a cross-sectional study. The Lancet 397, 1265–1275 (2021).

37. Scully, E. P., Haverfield, J., Ursin, R. L., Tannenbaum, C. & Klein, S. L. Considering how biological sex impacts immune responses and COVID-19 outcomes. Nat. Rev. Immunol. 20, 442–447 (2020).

38. Siam, M. H. B. et al. Insights into the first wave of the COVID-19 pandemic in Bangladesh: Lessons learned from a high-risk country. medRxiv 2020.08.05.20168674 (2020) doi:10.1101/2020.08.05.20168674.

39. Takahashi, T. et al. Sex differences in immune responses that underlie COVID-19 disease outcomes. Nature 588, 315–320 (2020).

40. Parvez, Md. S. A. et al. Genetic analysis of SARS-CoV-2 isolates collected from Bangladesh: Insights into the origin, mutational spectrum and possible pathomechanism. Comput. Biol. Chem. 90, 107413 (2021).

