## Supplementary Tables for "Investigating the possible origin and transmission routes of SARS-CoV-2 genomes and variants of concern in Bangladesh"

**Table S1**: Number of Samples for Initial Phylogeny from Different Countries and Territories (n=86,926)

| Region | Country / Territory | Sample Count |
| --- | --- | --- |
| Africa | Democratic Republic of the Congo (DRC) | 181 |
|  | Kenya | 94 |
|  | South Africa | 178 |
| Asia | Bangladesh | 612 |
|  | China | 830 |
|  | Hong Kong | 158 |
|  | India | 667 |
|  | Israel | 359 |
|  | Japan | 3,530 |
|  | Malaysia | 99 |
|  | Oman | 100 |
|  | Saudi Arabia | 523 |
|  | Singapore | 891 |
|  | South Korea | 239 |
|  | Taiwan | 122 |
|  | Thailand | 315 |
|  | United Arab Emirates (UAE) | 131 |
|  | Vietnam | 113 |
| Europe | Austria | 697 |
|  | Belgium | 854 |
|  | Denmark | 1,477 |
|  | Finland | 225 |
|  | France | 1,620 |
|  | Germany | 586 |
|  | Greece | 121 |
|  | Iceland | 577 |
|  | Ireland | 464 |
|  | Italy | 703 |
|  | Luxembourg | 258 |
|  | Netherlands | 1,786 |
|  | Norway | 89 |
|  | Poland | 125 |
|  | Portugal | 1471 |
|  | Russia | 314 |
|  | Slovenia | 107 |
|  | Spain | 3,975 |
|  | Sweden | 535 |
|  | Switzerland | 1571 |
|  | Turkey | 108 |
|  | United Kingdom | 30,822 |
| North America | Canada | 2,652 |
|  | Panama | 230 |
|  | USA | 20,944 |
| Oceania | Australia | 3,371 |
|  | New Zealand | 719 |
| South America | Brazil | 868 |
|  | Chile | 189 |
|  | Colombia | 160 |
|  | Peru | 166 |

**Table S2**: Number of B.1.1.7 Samples from Different Countries and Territories (n=7,507)

| Region | Country / Territory | Sample Count |
| --- | --- | --- |
| Africa | Democratic Republic of the Congo (DRC) | 7 |
|  | Egypt | 19 |
|  | Equatorial Guinea | 3 |
|  | Gambia | 42 |
|  | Ghana | 70 |
|  | Kenya | 133 |
|  | Morocco | 6 |
|  | Mozambique | 3 |
|  | Nigeria | 94 |
|  | Republic of the Congo | 2 |
|  | Rwanda | 4 |
|  | Senegal | 2 |
|  | South Africa | 119 |
|  | Tunisia | 4 |
|  | Zimbabwe | 2 |
| Asia | Bahrain | 1 |
|  | Bangladesh | 17 |
|  | Cambodia | 7 |
|  | China | 79 |
|  | Georgia | 2 |
|  | Hong Kong | 37 |
|  | India | 196 |
|  | Indonesia | 10 |
|  | Iran | 3 |
|  | Israel | 211 |
|  | Japan | 71 |
|  | Jordan | 44 |
|  | Kuwait | 3 |
|  | Malaysia | 6 |
|  | Myanmar | 1 |
|  | Nepal | 4 |
|  | Oman | 10 |
|  | Pakistan | 2 |
|  | Palestine | 1 |
|  | Philippines | 35 |
|  | Saudi Arabia | 28 |
|  | Singapore | 76 |
|  | South Korea | 31 |
|  | Sri Lanka | 25 |
|  | Taiwan | 5 |
|  | Thailand | 55 |
|  | United Arab Emirates | 28 |
|  | Vietnam | 4 |
| Europe | Austria | 148 |
|  | Belgium | 215 |
|  | Bosnia and Herzegovina | 5 |
|  | Croatia | 60 |
|  | Czech Republic | 17 |
|  | Denmark | 293 |
|  | Estonia | 1 |
|  | Finland | 94 |
|  | France | 224 |
|  | Germany | 136 |
|  | Greece | 4 |
|  | Hungary | 9 |
|  | Iceland | 27 |
|  | Ireland | 258 |
|  | Italy | 267 |
|  | Kosovo | 3 |
|  | Latvia | 5 |
|  | Lithuania | 1 |
|  | Luxembourg | 32 |
|  | Monaco | 1 |
|  | Montenegro | 7 |
|  | Netherlands | 272 |
|  | North Macedonia | 14 |
|  | Norway | 184 |
|  | Poland | 118 |
|  | Portugal | 232 |
|  | Romania | 61 |
|  | Russia | 1 |
|  | Serbia | 1 |
|  | Slovakia | 75 |
|  | Slovenia | 31 |
|  | Spain | 353 |
|  | Sweden | 170 |
|  | Switzerland | 254 |
|  | Turkey | 187 |
|  | Ukraine | 1 |
|  | United Kingdom | 552 |
| North America | Canada | 57 |
|  | Costa Rica | 2 |
|  | Dominican Republic | 1 |
|  | Guadeloupe | 4 |
|  | Mexico | 8 |
|  | Panama | 1 |
|  | USA | 594 |
| Oceania | Australia | 422 |
|  | New Zealand | 141 |
| South America | Argentina | 38 |
|  | Aruba | 54 |
|  | Bolivia | 2 |
|  | Brazil | 122 |
|  | Chile | 20 |
|  | Colombia | 86 |
|  | Curacao | 19 |
|  | Ecuador | 62 |
|  | Peru | 27 |
|  | Suriname | 9 |
|  | Trinidad and Tobago | 9 |
|  | Uruguay | 14 |

**Table S3**: Number of B.1.351 Samples from Different Countries and Territories (n=3,196)

| Region | Country / Territory | Sample Count |
| --- | --- | --- |
| Africa | Botswana | 19 |
|  | Egypt | 24 |
|  | France | 291 |
|  | Kenya | 33 |
|  | Mozambique | 57 |
|  | South Africa | 947 |
|  | Zambia | 31 |
| Asia | Bangladesh | 47 |
|  | China | 64 |
|  | Hong Kong | 10 |
|  | India | 11 |
|  | Israel | 17 |
|  | Japan | 26 |
|  | Singapore | 23 |
|  | United Arab Emirates | 9 |
| Europe | Austria | 105 |
|  | Belgium | 211 |
|  | Denmark | 17 |
|  | France | 124 |
|  | Italy | 12 |
|  | Netherlands | 90 |
|  | Switzerland | 75 |
|  | Turkey | 220 |
|  | United Kingdom | 229 |
| North America | Canada | 10 |
|  | USA | 163 |
| Oceania | Australia | 211 |
|  | New Zealand | 39 |
| South America | Brazil | 42 |
|  | Chile | 8 |
|  | Ecuador | 19 |
|  | Peru | 12 |

**Table S4:** Distribution of Age Group and Sex in G1 subset without missing values (n=836)

| Age Group | Male (n=551) | Female (n=285) |
| --- | --- | --- |
| < 10 | 48 | 20 |
| 10-19 | 18 | 18 |
| 20-29 | 71 | 64 |
| 30-39 | 136 | 49 |
| 40-49 | 111 | 57 |
| 50-59 | 95 | 44 |
| 60-69 | 48 | 25 |
| 70-79 | 19 | 6 |
| > 79 | 5 | 2 |

**Table S5:** Estimated Origins of Initial Representative Samples of Bangladesh

| Nextstrain Clade | Bangladeshi Sample | Estimated Origin |
| --- | --- | --- |
| 19A | EPI_ISL_458133 | United States |
| 19B | EPI_ISL_450339 | Australia |
|  | EPI_ISL_450341 |  |
|  | EPI_ISL_450345 | India |
| 20A | EPI_ISL_477134 |  |
|  | EPI_ISL_445244 | Italy |
|  | EPI_ISL_774877 | Germany |
|  | EPI_ISL_605887 |  |
|  | EPI_ISL_774896 |  |
|  | EPI_ISL_774900 | Russia |
|  | EPI_ISL_605910 |  |
|  | EPI_ISL_605889 | South Africa |
|  | EPI_ISL_468070 | Democratic Republic of the Congo |
|  | EPI_ISL_774893 |  |
|  | EPI_ISL_466637 | Saudi Arabia |
|  | EPI_ISL_450343 |  |
| 20B | EPI_ISL_477130 | United Kingdom |
|  | EPI_ISL_774890 |  |
|  | EPI_ISL_774889 |  |
|  | EPI_ISL_483630 | New Zealand |
|  | EPI_ISL_464162 |  |
|  | EPI_ISL_605884 | United Arab Emirates |
|  | EPI_ISL_514238 |  |
|  | EPI_ISL_774956 |  |
| 20C | EPI_ISL_477128 | Denmark |

**Table S6**: Estimated Origins of B.1.1.7 Variant Representative Samples of Bangladesh

| Nextstrain Clade | Bangladeshi Sample | Estimated Origin |
| --- | --- | --- |
| 20I/501Y.V1 | EPI_ISL_890237 | United Kingdom |
|  | EPI_ISL_1360439 |  |
|  | EPI_ISL_906091 | Spain |
|  | EPI_ISL_1508943 | Indonesia |
|  | EPI_ISL_1508946 |  |
|  | EPI_ISL_906098 | Ireland |
|  | EPI_ISL_1360445 | Australia |
|  | EPI_ISL_1498132 | India |
|  | EPI_ISL_1509000 | Nigeria |
|  | EPI_ISL_1360451 |  |
|  | EPI_ISL_1360430 |  |
|  | EPI_ISL_1360446 |  |
|  | EPI_ISL_1508954 |  |
|  | EPI_ISL_1508955 |  |

**Table S7:** Estimated Origins of B.1.351 Variant Representative Samples of Bangladesh

| Nextstrain Clade | Bangladeshi Sample | Estimated Origin |
| --- | --- | --- |
| 20H/501Y.V2 | EPI_ISL_1508892 | United States |
|  | EPI_ISL_943561 | South Africa |
|  | EPI_ISL_1360428 |  |
|  | EPI_ISL_1360447 |  |
|  | EPI_ISL_1360448 |  |
|  | EPI_ISL_1498150 |  |
